# Investigating low frequency somatic mutations in *Arabidopsis* with Duplex Sequencing

**DOI:** 10.1101/2024.01.31.578196

**Authors:** Gus Waneka, Braden Pate, J. Grey Monroe, Daniel B. Sloan

## Abstract

Mutations are the source of novel genetic diversity but can also lead to disease and maladaptation. The conventional view is that mutations occur randomly with respect to their environment-specific fitness consequences. However, intragenomic mutation rates can vary dramatically due to transcription coupled repair and based on local epigenomic modifications, which are non-uniformly distributed across genomes. One sequence feature associated with decreased mutation is higher expression level, which can vary depending on environmental cues. To understand whether the association between expression level and mutation rate creates a systematic relationship with environment-specific fitness effects, we perturbed expression through a heat treatment in *Arabidopsis thaliana*. We quantified gene expression to identify differentially expressed genes, which we then targeted for mutation detection using Duplex Sequencing. This approach provided a highly accurate measurement of the frequency of rare somatic mutations in vegetative plant tissues, which has been a recent source of uncertainty in plant mutation research. We included mutant lines lacking mismatch repair (MMR) and base excision repair (BER) capabilities to understand how repair mechanisms may drive biased mutation accumulation. We found wild type (WT) and BER mutant mutation frequencies to be very low (mean variant frequency 1.8×10^-8^ and 2.6×10^-8^, respectively), while MMR mutant frequencies were significantly elevated (1.13×10^-6^). These results show that somatic variant frequencies are extremely low in WT plants, indicating that larger datasets will be needed to address the fundamental evolutionary question as to whether environmental change leads to gene-specific changes in mutation rate.

**SIGNIFICANCE:** Accurately measuring mutations in plants grown under different environments is important for understanding the determinants of mutation rate variation across a genome. Given the low rate of *de novo* mutation in plant germlines, such measurements can take years to obtain, hindering tests of mutation accumulation under varying environmental conditions. We implemented highly accurate Duplex Sequencing to study somatic mutations in plants grown in two different temperatures. In contrast to plants with deficiencies in DNA mismatch repair machinery, we found extremely low mutation frequencies in wild type plants. These findings help resolve recent uncertainties about the somatic mutation rate in plant tissues and indicate that larger datasets will be necessary to understand the interaction between mutation and environment in plant genomes.

## INTRODUCTION

Mutations in DNA sequences accumulate over time and produce the variation that allows populations to adapt to novel or changing environments. In this sense, mutation is the ultimate source of evolutionary innovation. At the same time, mutations are often deleterious (Eyre-Walker and Keightley 2007), and somatic mutations can cause disease, setting up an interesting dynamic where selection may favor alleles that lower mutation rates, even though mutational input is required for adaptation and evolution (Zhang 2023).

The textbook view of mutation and adaptation is that mutations occur randomly with respect to their environment-specific fitness consequences. This principle was established in early investigations by Max Delbrück and Salvador Luria, who found that mutations in bacteria that confer phage resistance were equally likely to occur regardless of whether bacteria were grown in the presence of phage (Luria and Delbrück 1943). In other words, a phage-containing environment creates selection for genetic variants responsible for resistance but does not induce mutations to specifically occur at those loci. After subsequent decades of study, mutations are still widely considered to be random in this respect even though both the type and location of mutations are now known to have non-uniform distributions across genomes. For example, transition substitutions are far more common than transversions in most organisms across the tree of life. This bias in the mutation spectrum arises through the simple properties of DNA bases and chemical damage, but it has important consequences for the relationship between fitness effects and the probability of mutations. Due to the structure of the genetic code, transversions are more likely than transitions to be nonsynonymous (i.e. result in amino acid changes) and, therefore, have harmful fitness effects. As such, the average fitness effect of mutations is lower than it would be if all types of nucleotide substitutions occurred with equal probability (Eyre-Walker and Keightley 2007).

Mutation rates can also vary depending on genomic location. For example, mutational gradients arise in mammalian mitochondrial genomes because regions near replication origins are single-stranded (and more vulnerable to mutation causing damage) for longer periods during DNA replication (Sanchez-Contreras *et al*. 2021). Variation in intragenomic mutation rates can also occur at smaller scales, such is in *Arabidopsis thaliana* where mutations are enriched in intergenic sequences compared to genes (Ossowski *et al*. 2010; Belfield *et al*. 2018; Weng *et al*. 2019) and in introns compared to exons (Monroe *et al*. 2022, 2023a; Quiroz *et al*. 2023; Staunton *et al*. 2023). Because mutations in coding sequences are more likely to have functional consequences, this biased distribution of mutations should again result in lower average fitness effects than if mutations were uniformly distributed across the genome.

The probability of a mutation, therefore, cannot be considered independent of the fitness consequences of that mutation. However, to challenge the textbook view that mutations occur randomly with respect to environment-specific fitness effects, gene-specific mutational biases would have to systematically vary with changes in the environment. One potential mechanism that could create such a relationship between environment and mutation bias is the coupling of DNA repair surveillance with transcription machinery, which results in lower mutation rates for highly expressed genes (Supek and Lehner 2017; Oztas *et al*. 2018; Huang *et al*. 2018; Huang and Li 2018; Gonzalez-Perez *et al*. 2019; Monroe *et al*. 2022). Therefore, environmental changes that increase a gene’s expression level should lower its mutation rate. In addition, highly expressed genes are known to experience stronger selection (Zhang and Yang 2015), so genes may be most protected from mutation in environments where they are most functionally important. Alternatively, transcription may be mutagenic, as increased DNA damage associated with exposure of single-stranded DNA to mutagens can potentially overpower the increased protection of actively transcribed genes (Kim *et al*. 2007; Jinks-Robertson and Bhagwat 2014; Seplyarskiy *et al*. 2023).

A challenge associated with addressing how local mutation rates vary with environment is the difficulty of measuring mutations in experimental settings. Historical estimates of mutation relied on comparisons of synonymous substitutions between populations or species. Because these substitutions do not result in a change in amino acid, they are expected to experience minimal selection and thus approximate mutational input, though in reality synonymous sites do experience selection due to codon usage bias (Grantham *et al*. 1980; Hershberg and Petrov 2008) and other mechanisms (Bailey *et al*. 2021). It is inherently difficult to measure mutation rates more directly in large multicellular organisms because their long generations require many individuals and/or large amounts of time for sufficient mutations to occur, making methods such as mutation accumulation lines and parent-offspring trio sequencing (Lynch *et al*. 2016; Tatsumoto *et al*. 2017) expensive and time-consuming.

An alternative and potentially complementary approach to mutation accumulation and trio sequencing studies is to detect the mutations that accumulate in an organism’s somatic tissues (Gundry and Vijg 2012; Moore *et al*. 2021; Monroe *et al*. 2022; Quiroz *et al*. 2023; Schmitt *et al*. 2023; Staunton *et al*. 2023; Satake *et al*. 2023; Goel *et al*. 2024). This approach benefits from the fact that many more cell lineages can be tracked than just the germline. Inclusion of somatic (vegetative) mutations in recent *Arabidopsis* studies led to the identification of thousands of mutations, which increased power to test for relationships between local mutation rates and various sequence features, such as GC content, DNA methylation, histone modifications and expression level (Monroe *et al*. 2022). However, this approach appears to have been inaccurate because low frequency somatic variants can be difficult to distinguish from sequencing errors, and reanalysis of the somatic mutation calls showed that many of the putative mutations arose from technical artefacts (Liu and Zhang 2022; Monroe *et al*. 2023a; Wang *et al*. 2023; Monroe *et al*. 2023b). Therefore, the actual frequency of somatic mutations in vegetative plant tissue remains an open question.

Measurements of low frequency somatic mutations can be obtained using a high-fidelity sequencing technology to distinguish mutational signal from noise (Sloan *et al*. 2018). For example, Duplex Sequencing is an Illumina-based method in which unique molecular identifiers (UMIs) are included in adaptors and attached to both ends of DNA fragments before library amplification (Schmitt *et al*. 2012; Kennedy *et al*. 2014). After sequencing, the UMIs are used to cluster families of reads that originated from each strand of a given DNA fragment so that a double-stranded consensus sequence can be created that is virtually error free (< 5×10^-8^ errors per base pair; Kennedy *et al*. 2014).

Our goal in this study was to test if the pattern of local mutation rate variation across a genome depends on environmental effects on gene expression levels. We also wanted to determine whether low-frequency somatic mutations in plant tissues could provide a robust signal for addressing this type of question. Therefore, we perturbed gene expression by growing *Arabidopsis* under different temperatures. We identified differentially expressed (DE) genes with RNA-seq, which we then targeted for low-frequency somatic mutation detection using Duplex Sequencing coupled with hybrid capture. We included mutant lines *msh2* and *ung*, which respectively lack mismatch repair (MMR) and base excision repair (BER) capabilities, in order to understand how repair mechanisms may drive biased mutation accumulation (Cordoba-Canero *et al*. 2010; Belfield *et al*. 2018). We also included *hsp70-16* mutant lines, which are deficient for a key heat shock protein, as a means to endogenously manipulate gene expression and potentially interact with our temperature treatment (Ran *et al*. 2020). As expected, we found significant increases in variant frequencies in the MMR deficient lines. In wild type (WT) lines and other mutant lines, measured mutation frequencies were too low to quantify relationships between mutation rates and environment-specific gene expression levels. Therefore, our results support the conclusion that earlier estimates of somatic variant frequencies were inflated (Monroe *et al*. 2023a; Wang *et al*. 2023) and indicate that much larger datasets will be needed to test for environment-specific changes in mutation biases.

## RESULTS

To test if environment specific changes in gene expression impact mutation, we performed mutation detection on a targeted set of *Arabidopsis* genes that were DE in plants grown at 20°C vs. 30°C. We first generated and analyzed RNA-seq data to identify genes in six categories: 1) increased expression at 30°C compared to 20°C in WT plants, 2) increased expression at 20°C compared to 30°C in WT plants, 3) constitutively high expression in WT plants at both 20°C and 30°C, 4) constitutively low expression in WT plants at both 20°C and 30°C, 5) genes that had increased expression at 30°C vs. 20°C in WT plants (like category 1) and also had an interaction between WT and *hsp70-16*, and 6) genes that had increased expression at 30°C vs. 20°C in WT plants (like category 2) and also had an interaction between WT and *hsp70-16* (Table S1). The sequences of the DE genes were used to create a custom probe-set for hybrid capture of Duplex Sequencing libraries.

Duplex Sequencing coverage of the genes and 250 bp of flanking sequence in the probe-set ranged from 74.7× to 109.4× (Figure S1), and the average probe-set coverage across all libraries was 193.1-fold higher than the genome background. In total, we obtained 1.89 Gb of Duplex Sequencing coverage of our region of interest across the 24 libraries (Table S2)

We then looked for the presence of single nucleotide variants (SNVs) and short indels within the 339 genes covered in the probe-set. Mutant alleles already present in the parents of the assayed sets of full-sib plants have the potential to bias estimates of *de novo* mutation frequencies but should be readily identifiable. For a homozygous parent, they would be present in all Duplex Sequencing reads of all the replicates of a given genotype. For a heterozygous parent, they would segregate in a 1:2:1 Mendelian ratio and account for roughly 50% of the reads for all replicates of a given genotype (as each replicate represents a pool of five sibling plants). We identified just three apparent fixed SNVs (Table S3), which were removed for downstream analyses. In contrast, we identified 41 fixed indels, over half of which were in the *msh2* background (Table S4). One gene (AT5G39190) had five sites that appeared to be segregating SNVs in all 24 replicates. We suspected this might be caused by a cryptic gene duplication which was not captured in the TAIR 10.2 reference genome (Jaegle *et al*. 2023). Indeed, when we realigned the reads to the improved Col-CC genome (Reiser *et al*. 2023), the mutation calls in AT5G39190 were absent. As such, reads mapping to AT5G39190 were disregarded in downstream analyses. The rest of the SNVs we identified were unique to each replicate and all were present at a frequency of no more than 17.64% (the average variant frequency across all mutations was 2.27%), suggesting that these are low frequency somatic variants that arose during the experiment and were present in a subset of the sampled vegetative tissue.

Among the six WT biological replicates, we detected a single indel and just six SNVs, one in each replicate (Figure 1). As such, there was very limited statistical power to test for the effects of temperature or expression level on mutation frequency in WT plants. Similarly, we detected few or no SNVs and indels in the *hsp70-16* and the *ung* mutant lines (Figure 1; File S1, S2). In contrast, variant frequencies were significantly elevated in the *msh2* mutant lines (compared to WT plants), where we detected 271 indels and 180 SNVs (Figure 1; two-way ANOVA with Tukey’s test, *p* < 0.0001). The mutations in the *msh2* lines were distributed relatively evenly across the temperature treatments, as we found that temperature did not influence either SNV or indel frequency (Figure 1; two-way ANOVA, *p* = 0.99). In the *msh2* lines, deletions were 8.5-fold more common than insertions (Table S5; two-way ANOVA, *p* < 0.0001). We observed significant differences among SNV classes in *msh2* SNV spectrum (Figure 2; two-way ANOVA, *p* <0.0001), which was dominated by CG→TA transitions. The next most common types of substitutions were AT→GC transitions and CG→AT transversions. We compared the *msh2* mutation frequencies in the constitutively lowly expressed (group 3 in Table S1) vs constitutively highly expressed (group 4 in Table S1) genes and found no significant differences (paired t-test; Table S6), though we did observe a trend towards higher indel frequencies in constitutively highly expressed genes at 30°C. We did not analyze the SNV spectra or indel bias in WT, *ung*, *or hsp70-16* lines because the small number of sampled mutations precluded a statistically meaningful comparison.

**Figure 1.**
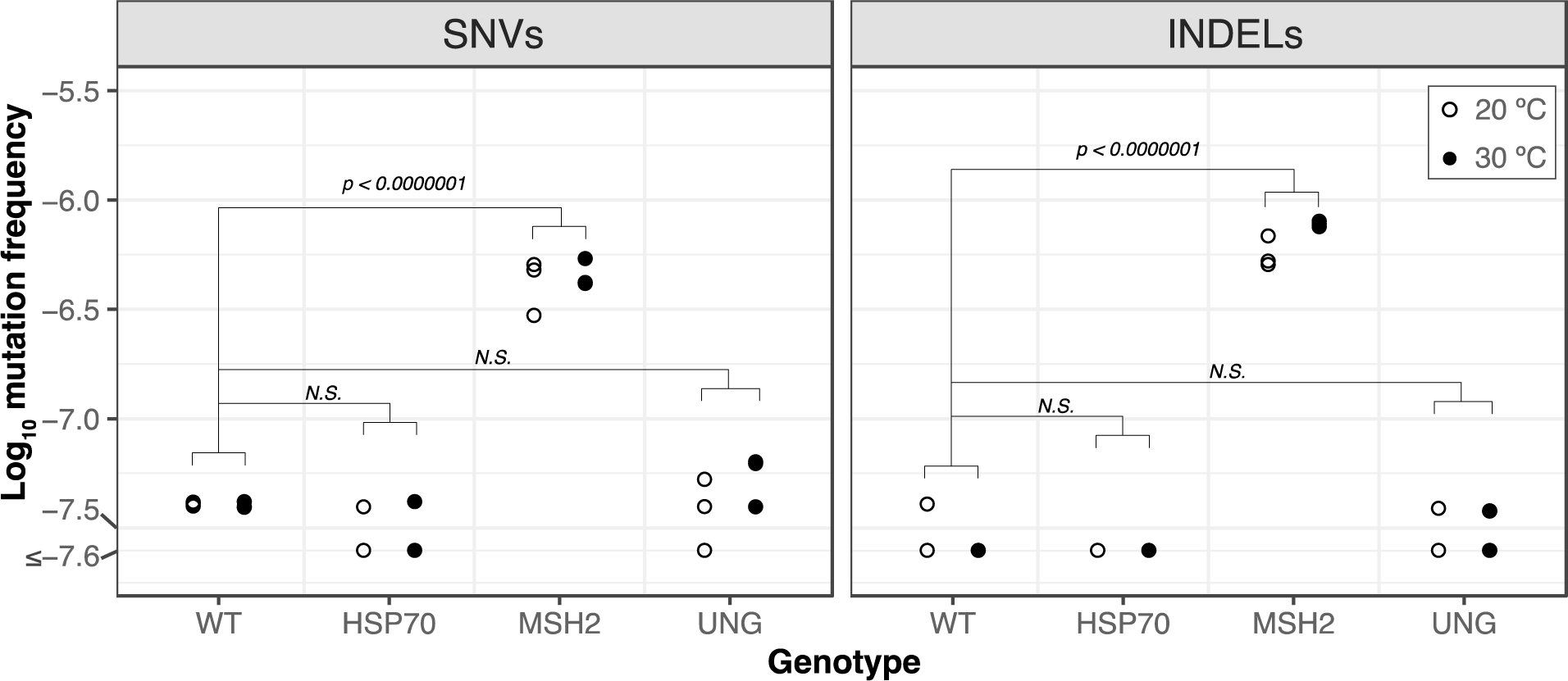
Mutation frequencies in WT vs mutant lines at 20°C and 30°C. Log_10_ mutation frequencies for single nucleotide variants (SNVs) and insertions/deletions (INDELs) calculated as the number of events (SNVs or INDELs) divided by the duplex sequencing coverage of the probe-set. A floor of 2.5×10^-8^ was applied to the y-axis for data visualization. *P-values* are from a Tukey’s test on a two-way ANOVA performed in R with the emmeans package (version 1; (Lenth *et al*. 2021).

**Figure 2.**
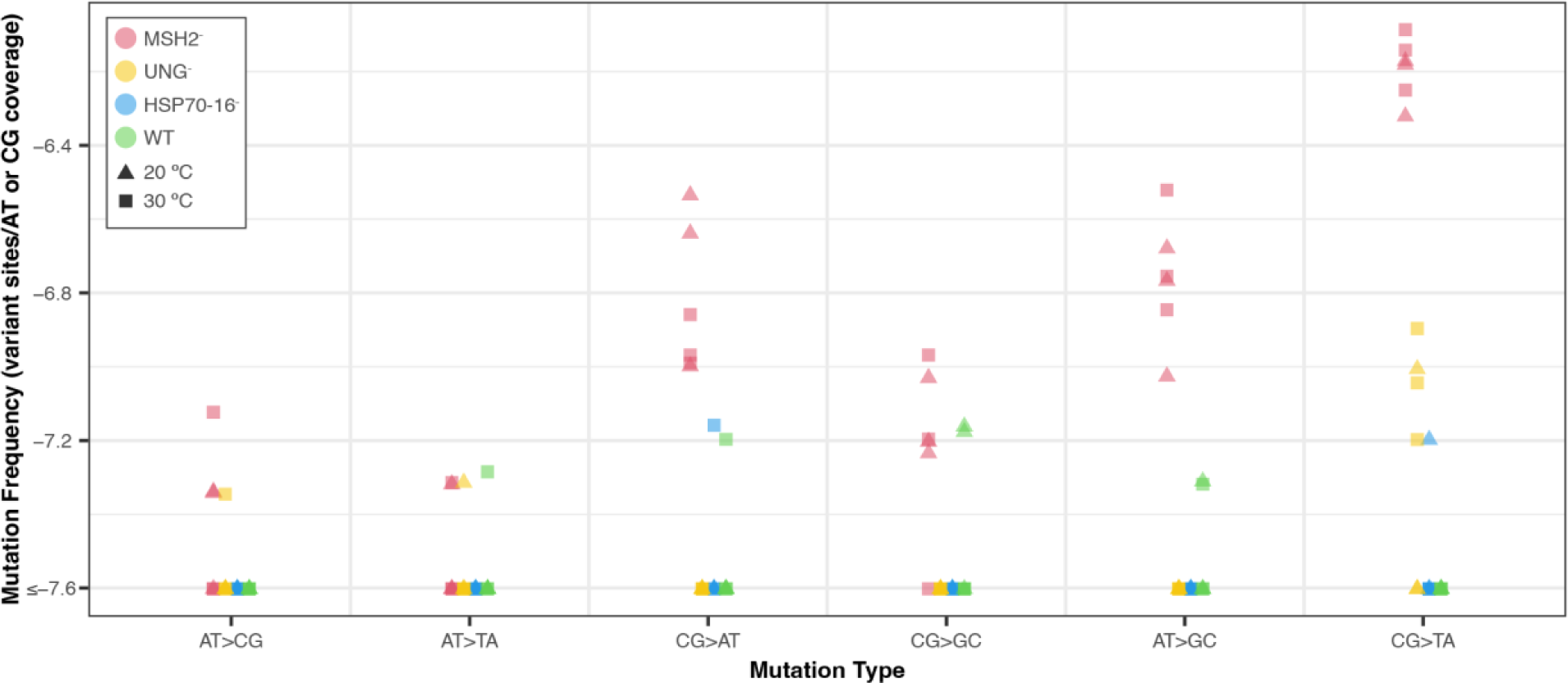
Mutation spectrum for WT and mutant plants at 20 °C and 30 °C. Log_10_ mutation frequencies for different types of single nucleotide variants were calculated as the number of events divided by the nucleotide-specific duplex sequencing coverage of the probe-set. A floor of 2.5×10^-8^ was applied to the y-axis for data visualization.

## DISCUSSION

In this study we took a novel approach to studying plant mutation by utilizing high fidelity Duplex Sequencing to measure low-frequency somatic variants in a targeted region of the *A. thaliana* nuclear genome. Variants in unopened floral bud tissue of WT plants were present at very low frequencies (Figure 1), which were near the detection threshold of Duplex Sequencing (Kennedy *et al*. 2014; Wu *et al*. 2020). Although we did not have enough power to address our prediction that increases in gene expression would correlate with decreases in mutation rates in WT plants, the results are nonetheless of interest given recent debates about the frequency of somatic mutations in plant tissues (Monroe *et al*. 2022; Liu and Zhang 2022; Monroe *et al*. 2023a; Wang *et al*. 2023; Monroe *et al*. 2023b). Our results support the conclusion that the high error rate of Illumina short-read sequencing makes it difficult to reliably discern sequencing errors from extremely rare WT somatic mutations. That said, we are skeptical of directly comparing the variant frequencies we measured in unopened floral buds with those obtained in differentiated leaves (Monroe *et al*. 2022, 2023a) given recent evidence showing substantial variation in somatic mutation rates depending on plant tissue (Goel *et al*. 2024).

We also surveyed variant frequencies in *ung* mutant plants and did not observe a difference between WT and *ung* lines. Given that *ung* plants have previously been shown to accumulate more uracil in DNA (presumably to the loss of base-excision repair activity on deaminated cytosines) than WT plants (Cordoba-Canero *et al*. 2010), we interpret the lack of a difference between WT and *ung* lines as evidence that actual WT mutation frequencies may be below the detection threshold of Duplex Sequencing. However, it is also possible that the similarly low mutation rates in WT and *ung* reflect the lack of a true biological difference, which may be possible if redundant pathways exist that prevent uracils in DNA from becoming CG→TA transitions.

In contrast, we found significantly elevated variant frequencies in *msh2* mutants compared to WT lines (Figure 1). MSH2 is known to function in mismatch repair (MMR) and mutation accumulation experiments with *msh2* mutant lines have established that the germline SNV rate is 132 to 204-fold greater than the WT SNV rate (Ossowski *et al*. 2010; Jiang *et al*. 2014; Belfield *et al*. 2018). Here, we found that the average *msh2* SNV frequency was 27-fold greater than the average WT SNV frequency (Figure 1). Though somatic variant frequencies measured with Duplex Sequencing are not directly comparable to germline mutation rates assayed with mutation accumulation experiments, the smaller magnitude of the difference between *msh2* vs. WT in our dataset may be interpreted as further evidence that the actual WT variant frequency is beneath the detection threshold of Duplex Sequencing. Alternatively, the smaller difference between WT and *msh2* reported here could be evidence that MMR is particularly important for buffering against mutation in germline plant tissues, which is supported by elevated expression of *MSH2* and other mismatch repair genes in meristematic tissues (Klepikova *et al*. 2016).

Variant frequencies in the *msh2* mutant lines showed no significant difference in plants grown at 20°C vs. 30°C. This finding contrasts with a recent mutation accumulation study that found elevated germline mutation rates in WT plants grown at 29°C compared to those grown at 23°C (Belfield *et al*. 2021) and another study that documented increases at 28°C and 32°C compared to 23°C (Lu *et al*. 2021). One potential explanation of this result is that heat stress may be mutagenic in WT plants *because* it impairs MMR since in the absence of MMR there is no apparent heat effect. However, this interpretation would be at odds with the fact that the genome-wide distribution of mutations in the heat-stressed plants mirrors the distribution of WT plants grown at standard temperature, not of mismatch repair mutants (see Figure 3 of (Belfield *et al*. 2021). The Duplex Sequencing variant frequencies in the *msh2* mutant lines also did not vary significantly between lowly expressed vs. highly expressed genes at either 20°C or 30°C (Figure 1). This result is consistent with the model that MMR provides special protection to actively transcribed genes (Belfield *et al*. 2018; Huang *et al*. 2018; Huang and Li 2018). However, we present this interpretation cautiously in the absence of WT data to test for an impact of expression when MMR is functional.

In summary, we took a novel approach to studying plant mutations by using Duplex Sequencing and hybrid capture to obtain a highly accurate snapshot of somatic variants in targeted regions of the *A. thaliana* genome. We designed our experiment to test if environmental conditions alter mutation rates in a gene-specific fashion. However, the low rate of mutations in WT plants prevented testing for how expression levels impact mutation rates. Nonetheless, the link between increased expression and decreased mutation in plants is well documented (Oztas *et al*. 2018; Monroe *et al*. 2022; Quiroz *et al*. 2023), as is the fact that gene expression is environmentally determined (Richards *et al*. 2012), so by logical extension environmental conditions must drive mutation rates and related fitness consequences. However, whether the magnitude of such an effect is biologically meaningful in shaping mutation and evolution remains an important, unanswered question. Though mutation accumulation and parent-offspring sequencing are time- and resource-intensive experiments, they are both increasingly feasible due to continued declines in the cost of DNA sequencing (Ossowski *et al*. 2010; Weng *et al*. 2019; Monroe *et al*. 2022). Conducting such experiments under contrasting environments (Jiang *et al*. 2014; Belfield *et al*. 2021; Lu *et al*. 2021) to measure the correlation between expression and mutation seems to be the key to understanding how environments impact the types of mutations that organisms accumulate.

## MATERIALS AND METHODS

All plants were grown in environmentally controlled growth chambers (75% humidity) under a long-day photoperiod (16 hrs light, 8 hrs dark) with irradiance of 185 µmol m^−2^ sec^−1^ at constant temperatures (either 20°C or 30°C, as specified below). Prior to planting, seeds were stratified for 5 days in sterile ddH20. *Arabidopsis thaliana* ecotype Col-0 was used as the WT line. Existing mutant lines were obtained from the Arabidopsis Biological Resource Center (Table S7) and seedlings were screened with allele-specific PCR markers to identify plants that were homozygous for the mutant alleles used in this study (*msh2*, *ung, hsp70-16;* Table S8).

Sibling plants (roughly 35 for each genotype and each temperature treatment) were planted in 2.5-inch pots. Both temperature treatments were initiated in chambers (Convarion models PGR15 (20°C) and PGCFLEX (30°C)) at 20°C because elevated ambient temperatures (30°C) can inhibit seed germination (Silva-Correia *et al*. 2014). After 5 days, the temperature was turned up for the 30°C treatment and kept at 20°C for the other treatment. When the plants had reached stage 6.5 of development (where ∼50 % of flowers have opened) (Boyes *et al*. 2001), we performed DNA and RNA extractions on unopened floral buds from laterally branching florets. The 30°C plants reached developmental stage 6.5 at 31 days while the 20°C plants reached developmental stage 6.5 at 41 days, consistent with faster plant development at elevated ambient temperatures (Silva-Correia *et al*. 2014).

For the RNA extractions, plant material was collected from the unopened floral buds of 3 laterally branching florets from 3 WT and 3 *hsp70-16* plants in each temperature treatment. The harvested tissues were immediately placed into liquid nitrogen and homogenized for 10 seconds at 30 beats/sec with the Qiagen TissueLyser, before being processed with the Qiagen RNeasy Plant Mini Kit, according to manufacturer’s instructions. The RNA samples were then sent to Novogene and RNA-Seq libraries were made using the NEBNext Ultra II Directional RNA Library Prep Kit with the NEBNext Poly(A) mRNA Magnetic Isolation Module. The RNA-Seq libraries were sequenced on a NovaSeq 6000 using the PE150 strategy to generate 29 to 54 million read pairs per library (see Table S9).

Tissue was harvested for DNA sequencing and mutation detection at the same time as the tissue for RNA extraction, from siblings of the plants used for RNA extraction. For each replicate in the DNA extractions, plant material was pooled from 5 siblings from the unopened floral buds of 3 laterally branching florets from 5 plants per each replicate, with 3 replicates per genotype (WT, *hsp70-16, msh2, ung*) per temperature treatment. The floret tissue was homogenized for 10 seconds at 30 beats/sec with the Qiagen TissueLyser, before being processed with the DNeasy Plant Mini Kit from Qiagen.

The RNA-seq reads were analyzed to detect DE genes at 20°C vs. 30°C. First, the adaptors were removed with Cutadapt version 4.0 with Python 3.9.16 (Martin 2011). Then the reads were mapped to the TAIR10.2 reference genome with HISAT2 (version 2.2.1; (Kim *et al*. 2019). Read counts were generated with HTSeq-count version 2.0.2 (Anders *et al*. 2014), and DESeq2 models (Love *et al*. 2014) were implemented to identify genes that were differentially expressed or constitutively highly or lowly expressed.

We created a custom probe-set to enrich the sequences of DE genes via hybrid capture so that we could perform mutation detection with Duplex Sequencing. We sent the sequences of 400 DE genes (plus 250 nt of flanking sequence on the end of each gene) to the probe design team at Arbor Bioscience, which flagged 61 of the genes as unsuitable for hybrid capture because they were > 25 % soft-masked for repeats in a BLAST search against the Arbor Biosciences eudicot database. The remaining 339 genes (listed in supplementary file 2) and flanking sequences spanned a total length of 855,123 nt. Sets of 80-nt probes were 2× tiled across the target sequence at approximately every 40 nt. The probes were biotinylated so that probe-bound library molecules can be captured with streptavidin-coated magnetic beads.

We created Duplex Sequencing libraries from the 24 DNA samples (3 replicates × 4 genotypes × 2 temperature treatments), following our previously described library preparation protocols (Wu *et al*. 2020; Waneka *et al*. 2021), except that in this case the amount of input DNA was increased to 500 ng because the target sequence comprises a small fraction (< 1%) of the total-cellular DNA sample. Once DNA samples had been fragmented via ultrasonication, end-repaired, A-tailed, adaptor-ligated, and treated with a cocktail of damage removal enzymes (Wu *et al*. 2020), we amplified 0.73 ng of DNA (per reaction) for 13 PCR cycles with New England Biolabs Q5 High-Fidelity Polymerase and dual-indexed primers. We then created 3 pools by combining 350 ng of each amplified library as the Arbor Biosciences hybrid-capture reactions have enough capacity for 8 libraries in each pool. We performed the overnight hybrid-capture reaction at 65°C, according to the manufacturer’s instructions (Arbor Biosciences MyBaits Kit Manual v. 5.02). We assessed enrichment efficiency and library concentrations through qPCR (as previously described; (Waneka *et al*. 2021)) before amplifying the enriched pools for an additional 9 cycles to obtain sufficient library amounts for sequencing.

Duplex Sequencing libraries were sequenced with PE150 reads on an Illumina NovaSeq 6000 S4 Lane (Novogene) to generate 87 to 123 million read pairs per library (Table S10). Processing of the Duplex Sequencing reads to was performed with our previously described pipeline (Wu *et al*. 2020), which trimmed adaptor sequences, created duplex consensus sequences based on the presence of shared barcodes, mapped the consensus sequences to the entire TAIR10.2 reference genome. Each duplex consensus sequences is composed of at least 6 Illumina reads (at least 3 originating from each strand of a DNA fragment). Alignment files were then parsed to identify duplex consensus sequences that contain SNVs and short indels. Since Duplex Sequencing is highly accurate (< 5×10^-8^ errors per base pair; Kennedy *et al*. 2014) we require just a single duplex consensus to support a putative mutation. Comparisons of coverage in the probe-set vs. outside the probe-set were performed with Samtools version 1.6 (Li *et al*. 2009). For variant frequency calculations, we excluded the first or last 10 bps of a read because we have previously identified elevated mutation frequencies at read ends (Wu *et al*. 2020).

## DATA AVAILABILITY

The raw reads are available via the NCBI Sequence Read Archive under accessions SRR27564102-SRR27564113 (RNA-seq libraries) and SRR27693810-SRR27693833 (Duplex Sequencing libraries). Duplex Sequencing datasets were processed with a previously published pipeline (https://github.com/dbsloan/duplexseq) (Wu *et al*. 2020).

## ACKNOWLEDGEMENTS

This work was supported by a grant from the National Institutes of Health (R35 GM148134).

## SUPPLEMENTARY FIGURES

**Figure S1.**
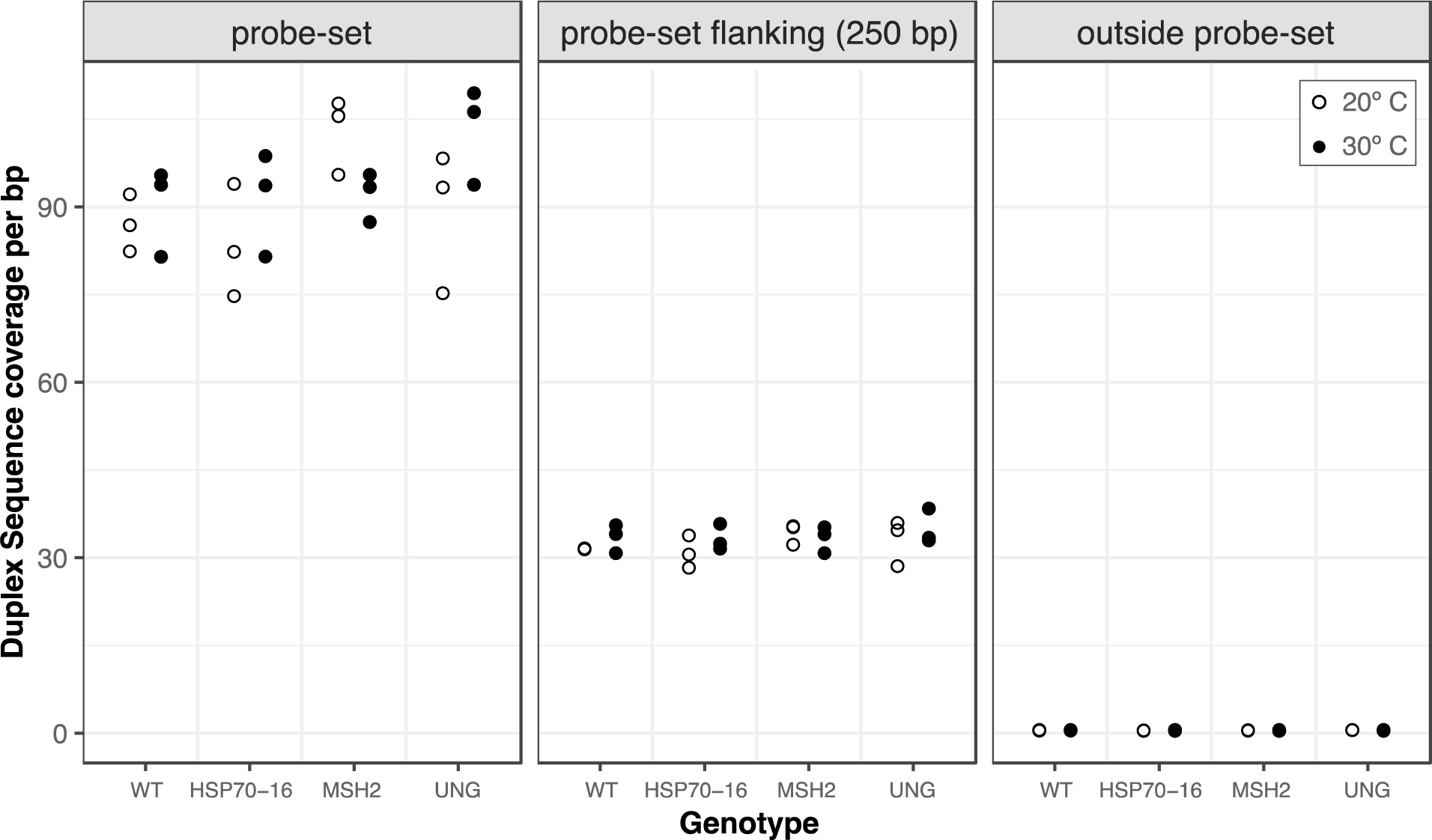
Duplex Sequencing coverage of the probe-set (panel 1), the 250 bps flanking the probe-set (panel 2) and the rest of the genome, outside of the probe-set (panel 3)

## SUPPLEMENTARY TABLES

**Table S1.** Differentially expressed genes from the RNA-seq analysis identified with DESeq2.

| Category | Genotype | Comparison | p-value | Log fold change | Average normalized cover of each treatment | Number of genes | Included in probe-set | Genes retained after arbor repeat filtering |
| --- | --- | --- | --- | --- | --- | --- | --- | --- |
| 1 | WT | Increased exp. at 30°C | 0.05 | > 2 | Minimum coverage > 5 | 683 | 100 with greatest LFC | 84 |
| 2 | WT | Increased exp. at 20°C | 0.05 | < -2 | Minimum coverage > 5 | 350 | 100 with lowest LFC | 80 |
| 3 | WT | Constitutive low exp. | 0.05 | 50 genes with LFC closest to 0 | 50 genes with lowest coverage (ranges from 129 to 400) | 50 | 50 | 44 |
| 4 | WT | Constitutive high exp. | 0.05 | 50 genes with LFC closest to 0 | 50 genes with highest coverage (ranges from 8384 to 68053) | 50 | 50 | 45 |
| 5 | WT vs. HSP70-16 | Interaction between genotype and temp | 0.05 | >2 | Minimum coverage > 5 | 106 (39 of which are also in group 1) | 92 with highest LFC | 81 |
| 6 | WT vs HSP70-60 | Interaction between genotype and temp | 0.05 | <-2 | Minimum coverage > 5 | 8 (5 of which are also in group 2) | All 8 | 5 |
| total |  |  |  |  |  |  | 400 | 339 |

**Table S2.**
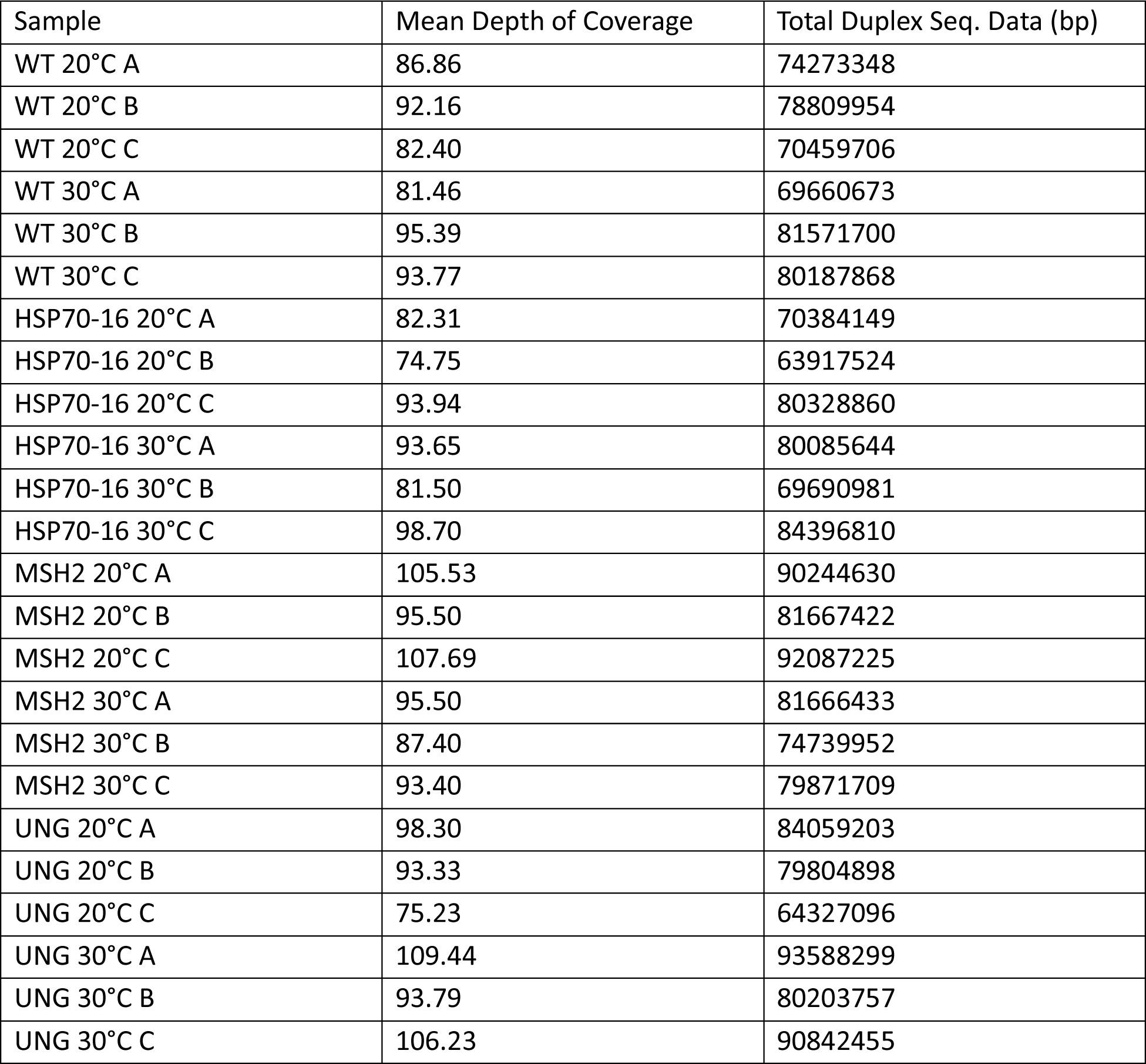
Duplex Sequencing coverage for each replicate.

**Table S3.** Putative fixed SNVs removed before downstream analysis of Duplex Sequencing data.

| Genotype | Chromosome | Position | Substitution type | Shared among all replicates |
| --- | --- | --- | --- | --- |
| *ung* | 2 | 2016156 | AT→GC | yes |
| *wild-type* | 2 | 14827204 | CG→AT | yes |
| *msh2* | 4 | 14827204 | CG→AT | yes |

**Table S4.**
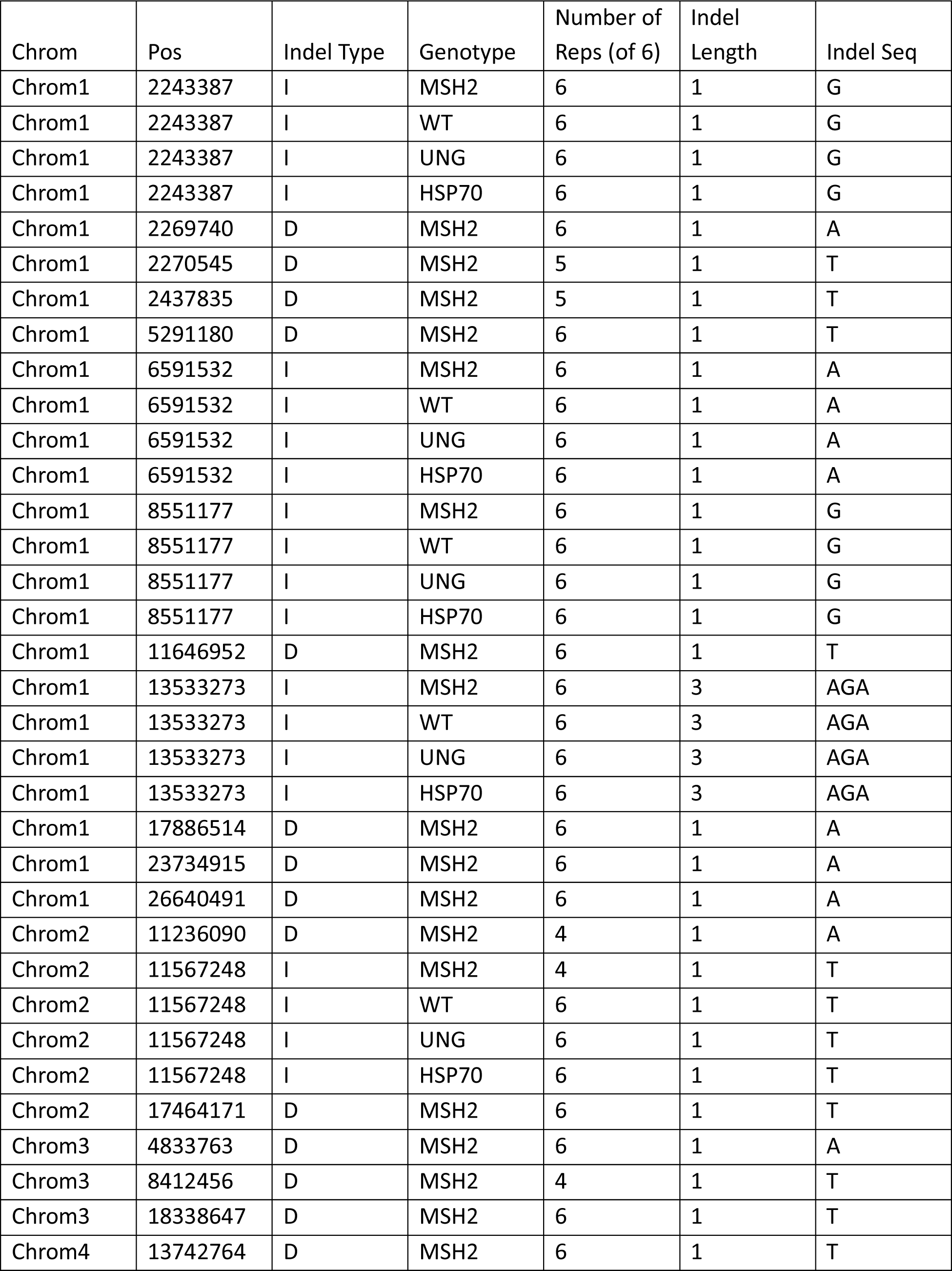
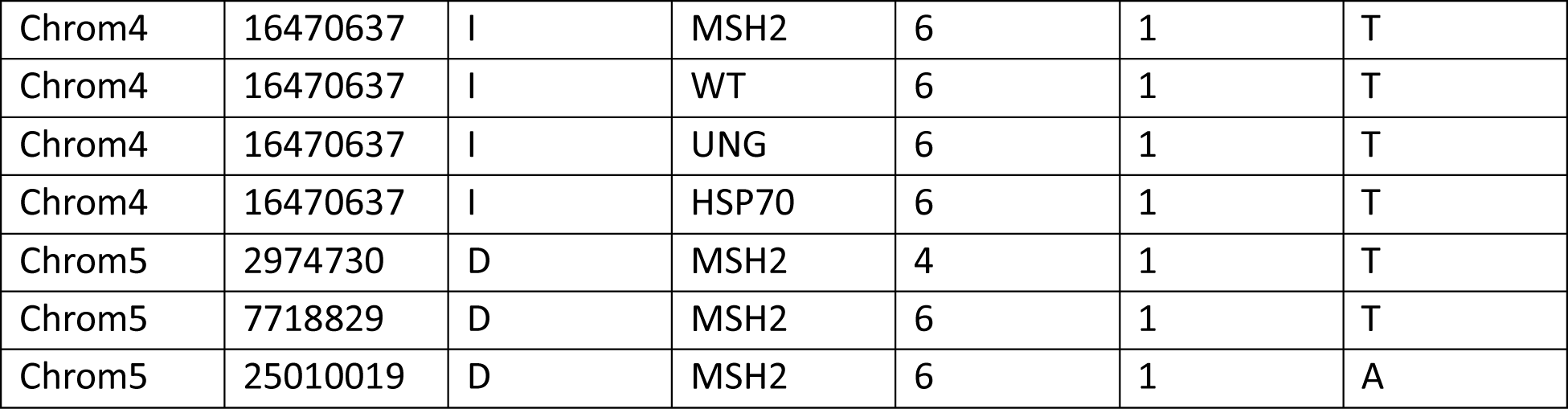
Putative fixed indels removed before downstream analysis of Duplex Sequencing data.

| Chrom | Pos | Indel Type | Genotype | Number of<br>Reps (of 6) | Indel<br>Length | Indel Seq |
| --- | --- | --- | --- | --- | --- | --- |
| Chrom1 | 2243387 | I | MSH2 | 6 | 1 | G |
| Chrom1 | 2243387 | I | WT | 6 | 1 | G |
| Chrom1 | 2243387 | I | UNG | 6 | 1 | G |
| Chrom1 | 2243387 | I | HSP70 | 6 | 1 | G |
| Chrom1 | 2269740 | D | MSH2 | 6 | 1 | A |
| Chrom1 | 2270545 | D | MSH2 | 5 | 1 | T |
| Chrom1 | 2437835 | D | MSH2 | 5 | 1 | T |
| Chrom1 | 5291180 | D | MSH2 | 6 | 1 | T |
| Chrom1 | 6591532 | I | MSH2 | 6 | 1 | A |
| Chrom1 | 6591532 | I | WT | 6 | 1 | A |
| Chrom1 | 6591532 | I | UNG | 6 | 1 | A |
| Chrom1 | 6591532 | I | HSP70 | 6 | 1 | A |
| Chrom1 | 8551177 | I | MSH2 | 6 | 1 | G |
| Chrom1 | 8551177 | I | WT | 6 | 1 | G |
| Chrom1 | 8551177 | I | UNG | 6 | 1 | G |
| Chrom1 | 8551177 | I | HSP70 | 6 | 1 | G |
| Chrom1 | 11646952 | D | MSH2 | 6 | 1 | T |
| Chrom1 | 13533273 | I | MSH2 | 6 | 3 | AGA |
| Chrom1 | 13533273 | I | WT | 6 | 3 | AGA |
| Chrom1 | 13533273 | I | UNG | 6 | 3 | AGA |
| Chrom1 | 13533273 | I | HSP70 | 6 | 3 | AGA |
| Chrom1 | 17886514 | D | MSH2 | 6 | 1 | A |
| Chrom1 | 23734915 | D | MSH2 | 6 | 1 | A |
| Chrom1 | 26640491 | D | MSH2 | 6 | 1 | A |
| Chrom2 | 11236090 | D | MSH2 | 4 | 1 | A |
| Chrom2 | 11567248 | I | MSH2 | 4 | 1 | T |
| Chrom2 | 11567248 | I | WT | 6 | 1 | T |
| Chrom2 | 11567248 | I | UNG | 6 | 1 | T |
| Chrom2 | 11567248 | I | HSP70 | 6 | 1 | T |
| Chrom2 | 17464171 | D | MSH2 | 6 | 1 | T |
| Chrom3 | 4833763 | D | MSH2 | 6 | 1 | A |
| Chrom3 | 8412456 | D | MSH2 | 4 | 1 | T |
| Chrom3 | 18338647 | D | MSH2 | 6 | 1 | T |
| Chrom4 | 13742764 | D | MSH2 | 6 | 1 | T |
| Chrom4 | 16470637 | I | MSH2 | 6 | 1 | T |
| Chrom4 | 16470637 | I | WT | 6 | 1 | T |
| Chrom4 | 16470637 | I | UNG | 6 | 1 | T |
| Chrom4 | 16470637 | I | HSP70 | 6 | 1 | T |
| Chrom5 | 2974730 | D | MSH2 | 4 | 1 | T |
| Chrom5 | 7718829 | D | MSH2 | 6 | 1 | T |
| Chrom5 | 25010019 | D | MSH2 | 6 | 1 | A |

**Table S5.**
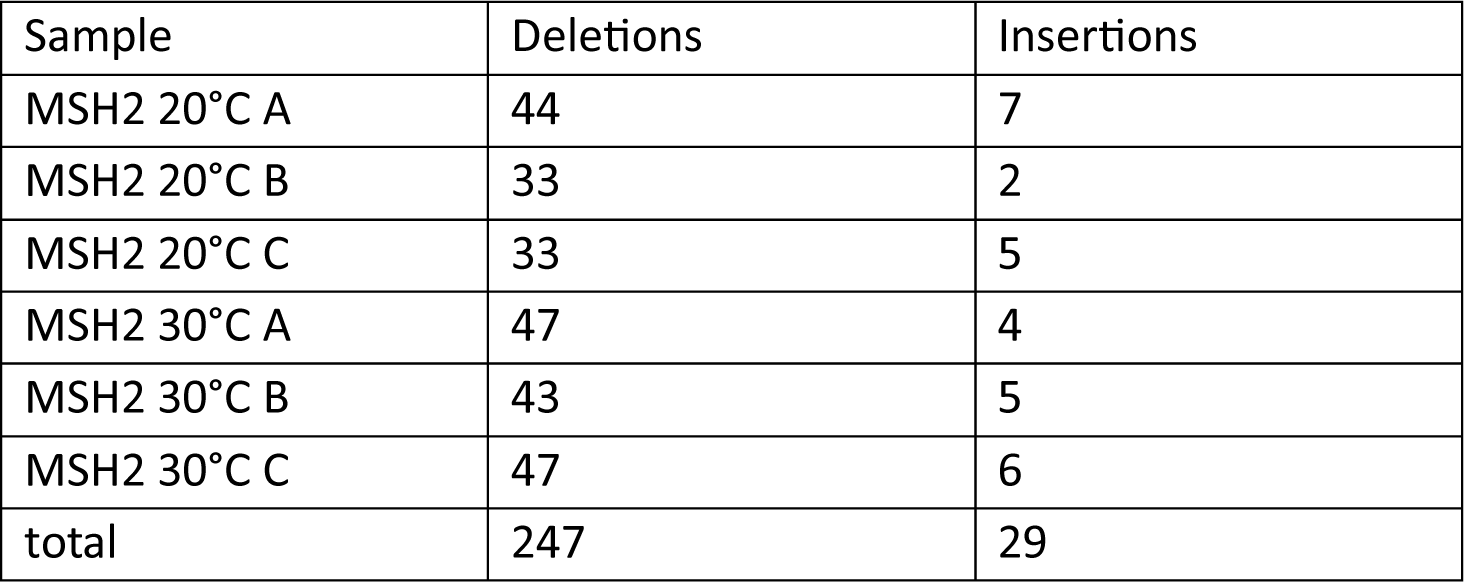
Indel mutations in *msh2* mutant lines.

| Sample | Deletions | Insertions |
| --- | --- | --- |
| MSH2 20°C A | 44 | 7 |
| MSH2 20°C B | 33 | 2 |
| MSH2 20°C C | 33 | 5 |
| MSH2 30°C A | 47 | 4 |
| MSH2 30°C B | 43 | 5 |
| MSH2 30°C C | 47 | 6 |
| total | 247 | 29 |

**Table S6.**
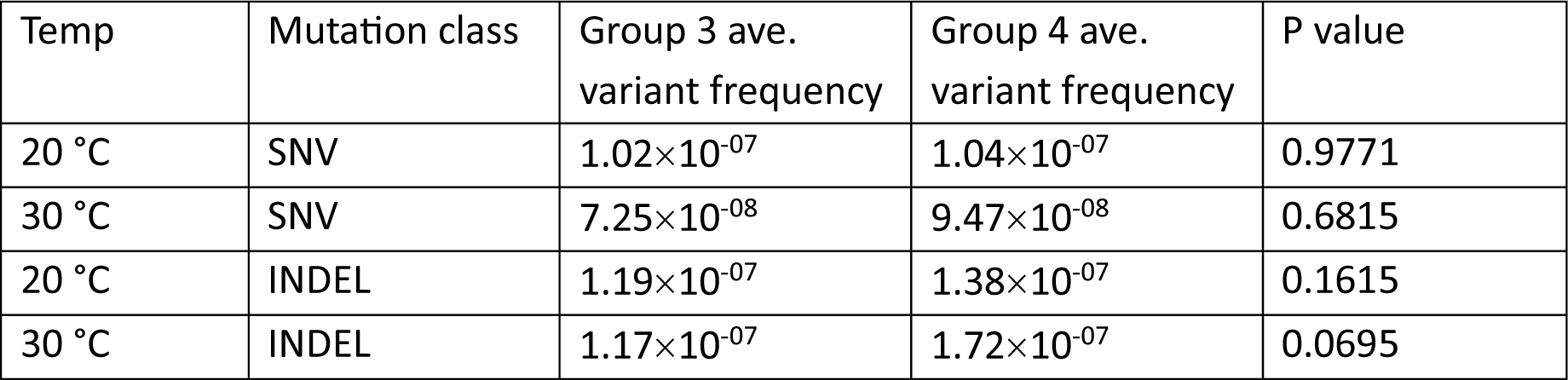
Paired t-test results of group 3 vs group 4 mutation rates in *msh2^−^* lines (two-tailed)

| Temp | Mutation class | Group 3 ave.<br>variant frequency | Group 4 ave.<br>variant frequency | P value |
| --- | --- | --- | --- | --- |
| 20 °C | SNV | $1.02 \times 10^{-07}$ | $1.04 \times 10^{-07}$ | 0.9771 |
| 30 °C | SNV | $7.25 \times 10^{-08}$ | $9.47 \times 10^{-08}$ | 0.6815 |
| 20 °C | INDEL | $1.19 \times 10^{-07}$ | $1.38 \times 10^{-07}$ | 0.1615 |
| 30 °C | INDEL | $1.17 \times 10^{-07}$ | $1.72 \times 10^{-07}$ | 0.0695 |

**Table S7.** Mutant lines used, all sourced from ABRC.

| Gene | AGI | Mutant Allele | Ref |
| --- | --- | --- | --- |
| HSP70-16 | AT1G11660 | SALK_028829 | (Ran *et al.* 2020) |
| MSH2 | AT3G18524 | SALK_002708 | (Belfield *et al.* 2018) |
| UNG | AT3G18630 | CS308297 | (Cordoba-Canero *et al.* 2010) |

**Table S8.**
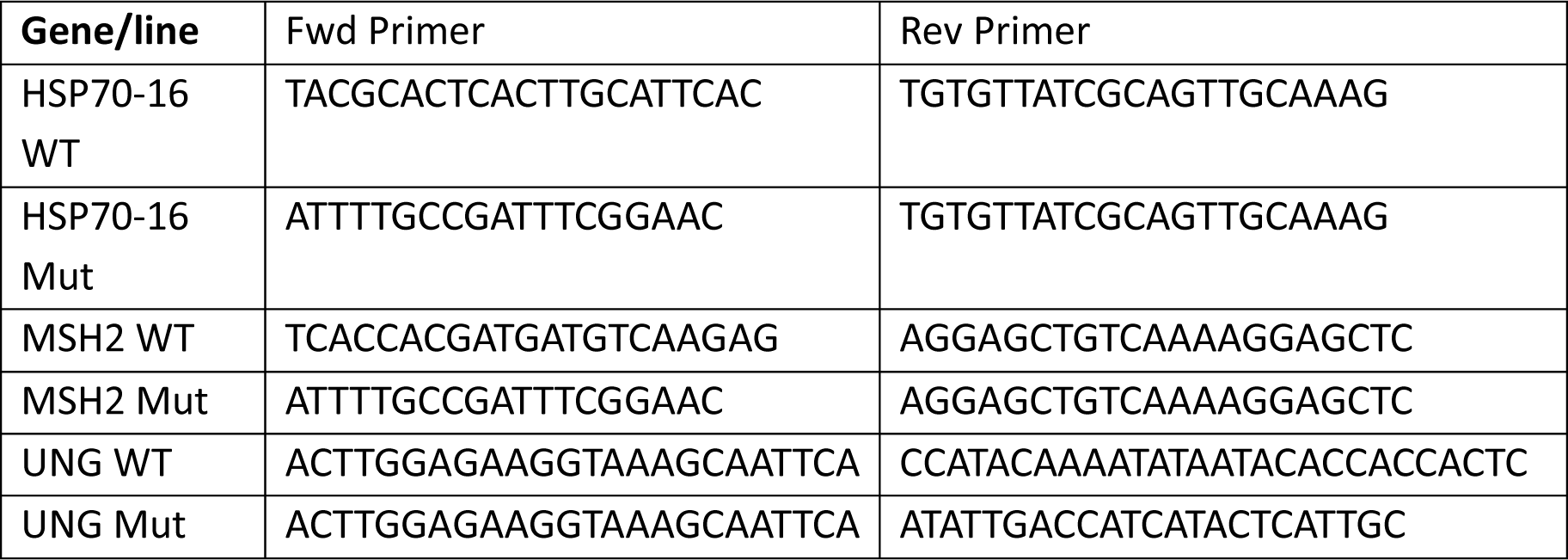
PCR primers used to identify mutant alleles in the three mutant lines.

**Table S9.** Read counts for the 12 RNA-seq libraries.

| Sample | Count of read pairs |
| --- | --- |
| HSP70-16 20°C A | 29689895 |
| HSP70-16 20°C B | 32052311 |
| HSP70-16 20°C C | 33450418 |
| HSP70-16 30°C A | 32567642 |
| HSP70-16 30°C B | 31456737 |
| HSP70-16 30°C C | 29678098 |
| WT 20°C A | 30417658 |
| WT 20°C B | 54410188 |
| WT 20°C C | 42449872 |
| WT 30°C A | 34353207 |
| WT 30°C B | 36605678 |
| WT 30°C C | 37953073 |

**Table S10.**
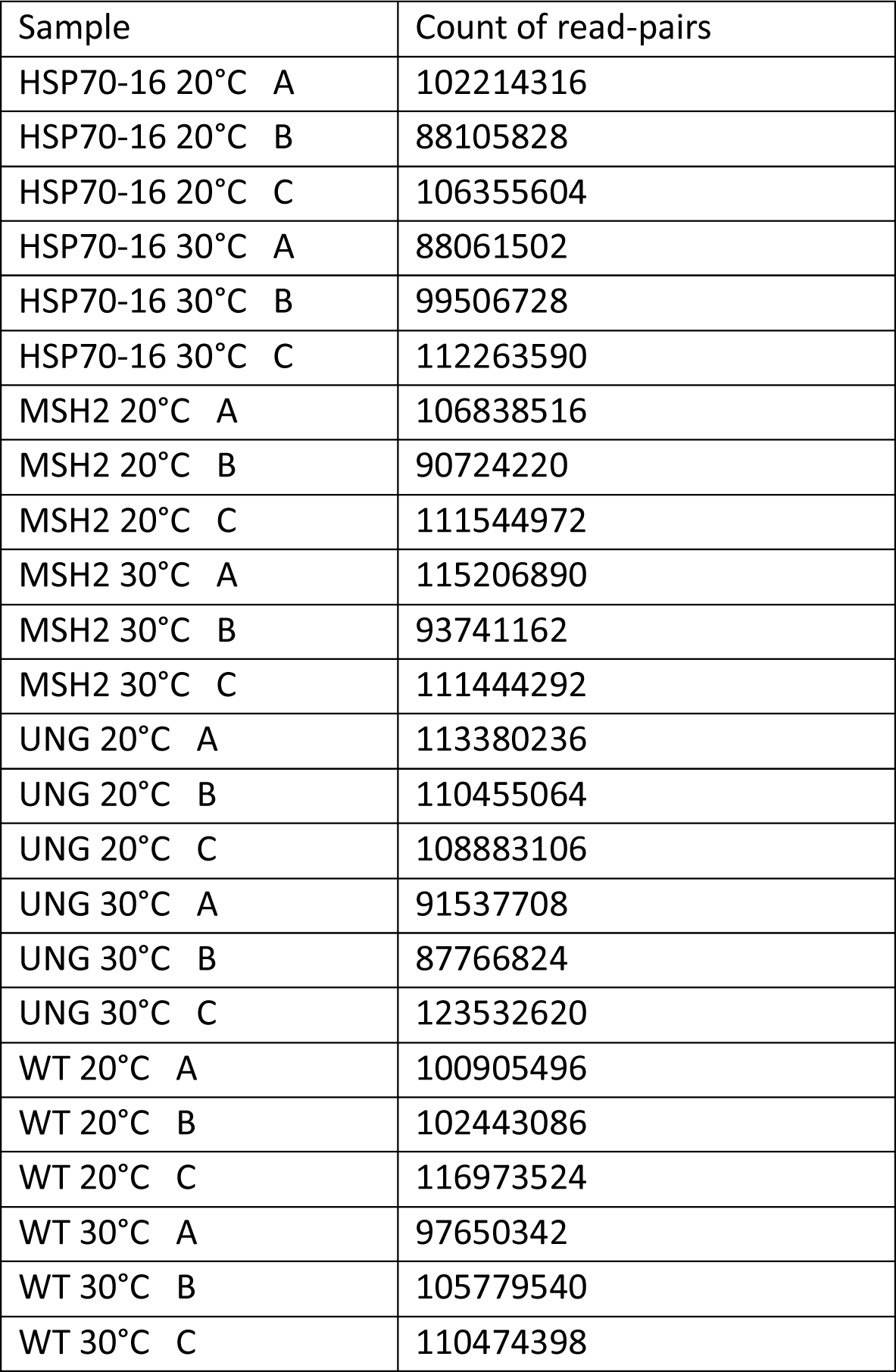
Read counts for the 24 Duplex Sequencing libraries.

| Sample | Count of read-pairs |
| --- | --- |
| HSP70-16 20°C A | 102214316 |
| HSP70-16 20°C B | 88105828 |
| HSP70-16 20°C C | 106355604 |
| HSP70-16 30°C A | 88061502 |
| HSP70-16 30°C B | 99506728 |
| HSP70-16 30°C C | 112263590 |
| MSH2 20°C A | 106838516 |
| MSH2 20°C B | 90724220 |
| MSH2 20°C C | 111544972 |
| MSH2 30°C A | 115206890 |
| MSH2 30°C B | 93741162 |
| MSH2 30°C C | 111444292 |
| UNG 20°C A | 113380236 |
| UNG 20°C B | 110455064 |
| UNG 20°C C | 108883106 |
| UNG 30°C A | 91537708 |
| UNG 30°C B | 87766824 |
| UNG 30°C C | 123532620 |
| WT 20°C A | 100905496 |
| WT 20°C B | 102443086 |
| WT 20°C C | 116973524 |
| WT 30°C A | 97650342 |
| WT 30°C B | 105779540 |
| WT 30°C C | 110474398 |

